# TraceQC: An R package for quality control of CRISPR lineage tracing data

**DOI:** 10.1101/2020.06.05.137141

**Authors:** Jingyuan Hu, Rami Al-Ouran, Xiang Zhang, Zhandong Liu, Hyun-Hwan Jeong

## Abstract

**Motivation:** The CRISPR-based lineage tracing system is emerging as a powerful new sequencing tool to track cell lineages by marking cells with irreversible genetic mutations. Accurate reconstruction of cell lineages from CRISPR-based data is sensitive to noise. Quality control is critical for filtering out low-quality data points. Yet, existing quality control tools for RNA-seq and DNA-seq do not measure features specific for the CRISPR-linear tracing system.

**Results:** We introduce TraceQC, a quality control package to overcome challenges with measuring the quality of CRISPR-based lineage tracing data and help in interpreting and constructing lineage trees.

**Availability:** The R package is available at https://github.com/LiuzLab/TraceQC.

## 1. Introduction

Determining the origin of cells has been a challenging quest in the field of developmental biology, for which lineage tracing methods have been used to identify and track cell progeny (Wu *et al*., 2019; Kretzschmar and Watt, 2012). The traditional lineage tracing techniques such as Cre-Lox recombination has limitations in the number of cells marked per experiment. Recently, a new generation of lineage tracing tools using CRISPR technology have been developed to track massive cells in a single experiment (McKenna *et al*., 2016). CRISPR-based lineage tracing system can mark cell with irreversible genetic mutations, thus simultaneously recording cell lineage along with its development.

There are three major challenges in reconstructing lineage from this new type of CRISPR data. First, CRISPR-induced mutations are not uniformly distributed (Chen *et al*., 2019). Lineage tracing assumes cells that share the same mutation event arise from the same lineage. However, some dominant mutation patterns may not be inherited but rather arise independently from mutation hotspot.

Second, the doxycycline-regulated inducible CRISPR system(Kalhor *et al*., 2017), which is commonly used in lineage tracing, is known for its leakiness of Cas9 expression(Dow *et al*., 2015). As a result, the lineage tracing system could be contaminated with many background mutations before the induction of Doxycycline. Therefore, it is critical to measure the level of background mutation before the induction for accurate lineage tracing.

Third, (Salvador-Martinez *et al*., 2019) found that the mutation rate (number of CRISPR mutations per cell division) has a strong impact on the accuracy of the lineage tree reconstruction. The study has also shown that mutation rate should be kept within a certain range for optimal results. A low mutation rate will not generate enough signals per cell cycle while a high mutation rate will cause the system to converge quickly. Therefore, measuring the mutation rate is key to understand the capacity of the CRISPR system in tracking individual cells.

To address the above challenges, we developed TraceQC, a quality control package to evaluate the quality of CRISPR lineage tracing data. TraceQC provides various functions to assist users to interpret the distribution, leakiness level, and rate of mutations of CRISPR-based lineage tracing data.

## 2 Methods

The complete workflow of TraceQC is shown in Supplementary Figure 1. Each subsection below describes a module of the TraceQC report.

**Fig. 1.**
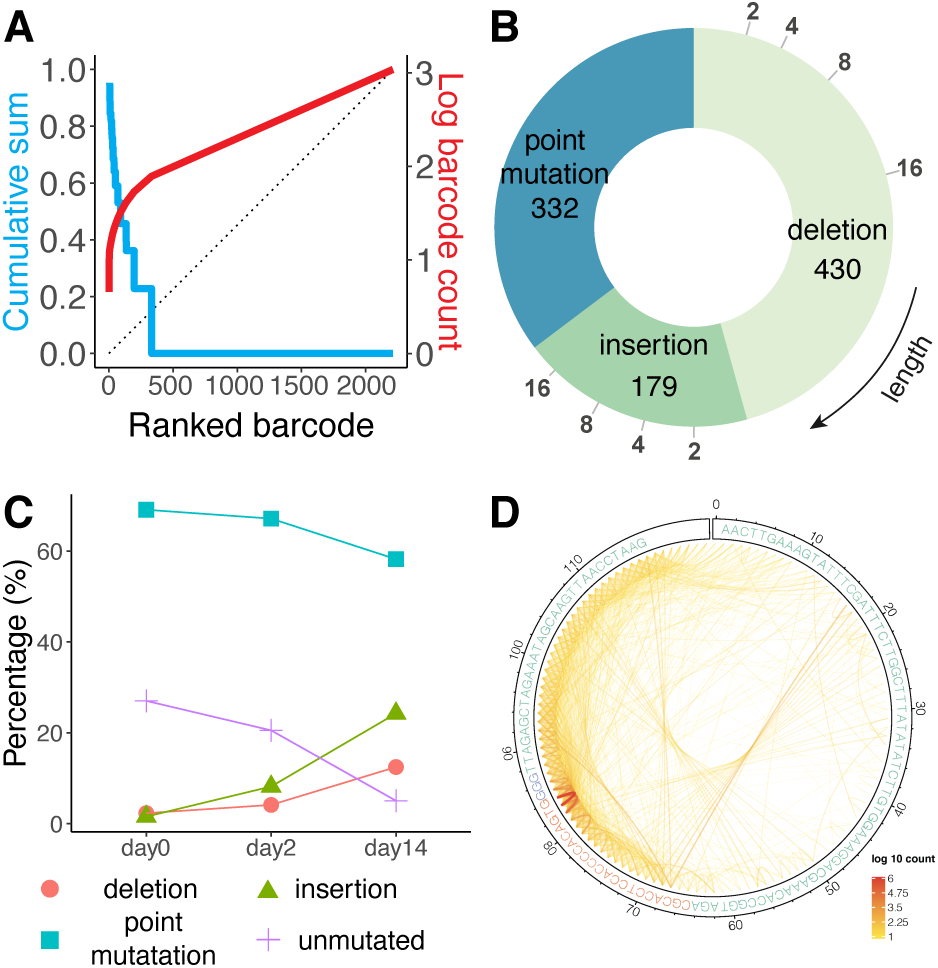
Examples of TraceQC output. (A) The Lorenz curve ranks the barcode from the most frequent one to the least, suggesting barcodes are not uniformly distributed. (B) The pie chart shows the mutation diversity of three main mutation types (insertion, deletion, and point mutation). The indel mutations are sorted by length as shown on the x axis. (C) The mutation rate can be inferred using samples at different time points. Day0 sample is sequenced before Cas9 induction. (D) The chord diagram shows the mutation hotspots.

### 2.1 Estimation of Barcode Distribution by Lorenz curve

A DNA barcode is a DNA sequence where CRISPR targeted mutation occurs. A Lorenz curve (Lorenz, 1905) indicates that the barcode distribution is not uniform (Figure 1A). Over-represented barcodes result from one or more of the following: 1) Clonal expansion of mutated barcode, 2) CRISPR mutation hotspot. This QC measure will allow users to detect any over-represented barcodes and estimate the barcode distribution of the input data.

### 2.2 Summary of Mutation Diversity

After mutation events are extracted from each barcode (Supplementary Method), TraceQC summarizes the mutation diversity in terms of insertion, deletion, and point mutation (Figure 1B). Since CRISPR-induced mutations are dominated by indels (Chen *et al*., 2019), TraceQC provides an easy visualization aid to help users identify any abnormality in the frequency and length distribution of indel events. Moreover, TraceQC provides another visualization on the number of mutations per sequence (Supplementary Method), which allows users to inspect whether recurrences of CRISPR-induced mutations had happened.

### 2.3 Evaluation of System Leakiness and Mutation Rate

Leakiness is a known issue in an inducible CRISPR system, in which the Cas9 protein expression is activated before induction. To evaluate the level of system leakiness, TraceQC calculates the background mutation level by examining the input before the induction starts. When combined with mutation levels from other time points, users can examine the capacity of the CRISPR system in tracking individual cells (Figure 1C).

### 2.4 Detection of Mutation Hotspot

As mentioned before, CRISPR-induced mutations are not uniformly distributed, and certain dominant mutations have been observed across multiple samples (Kalhor *et al*., 2017). These mutations arise independently during non-homologous end-joining rather than being inherited from common ancestors. Our package provides visualization on CRISPR-induced mutational hotspots (Figure 1D), which allows users to select genuine mutation patterns by region and frequency.

## 3 Discussion and Conclusion

We developed TraceQC to examine the quality of CRISPR-based lineage data. TraceQC can assist users to inspect and filter lineage data, and as a result, accurately construct the lineage tree. In addition to providing quality control for CRISPR-based lineage tracing data, TraceQC can transform raw sequencing data to a list of Boolean sequences of mutation events (Supplementary method) so that users do not need any pre-processing of the raw data to run their downstream analysis (Supplementary Method).

In conclusion, CRISPR-based lineage tracing technology leads a new generation of tools that can trace cell lineage at single cell resolution. TraceQC provides a series of modules of quality control measures on lineage tracing data including divergence, distribution, and rate of mutation. TraceQC is open-source and provides easy-to-use functions to examine the quality of the CRIPSR lineage tracing data.

## Acknowledgements

**Funding**

This work has been supported by CPRIT RP170387, Hamill and Chao Endowment.

