## Supplementary Material for "TraceQC: An R package for quality control of CRISPR lineage tracing data"

6/3/2020

#### Supplementary Figures

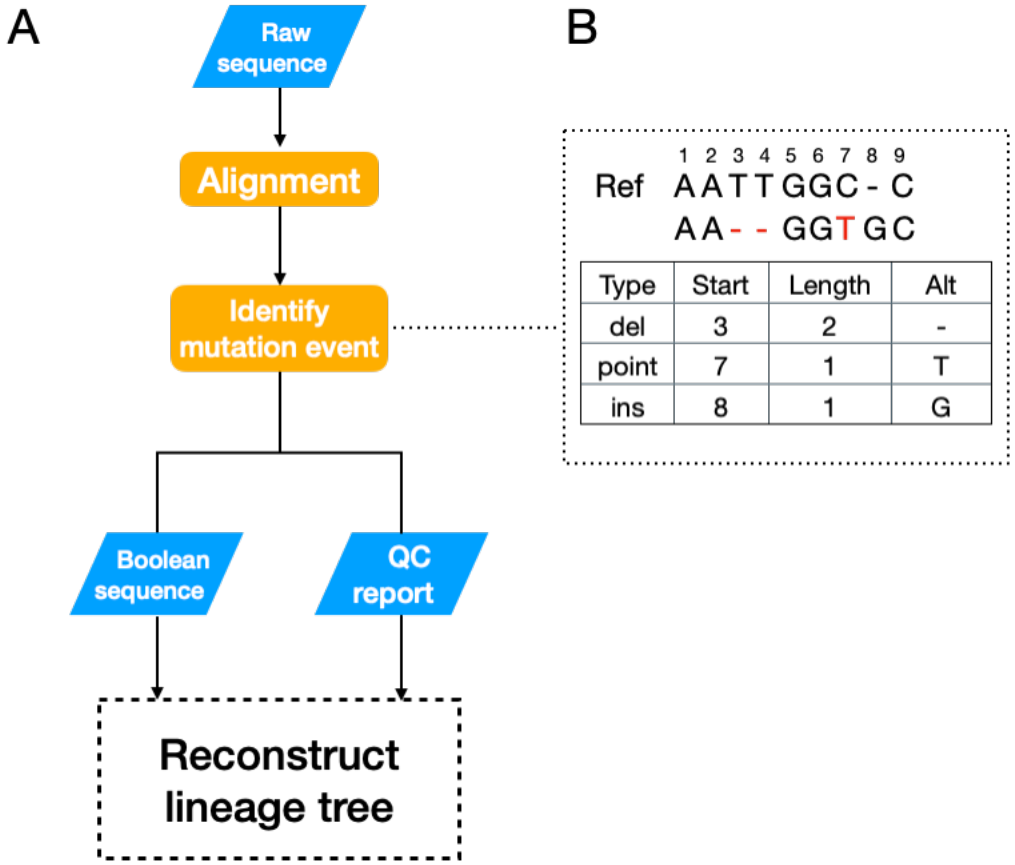

Figure S1: (A) Workflow of using TraceQC as a data analysis pipeline. (B) TraceQC identifies each mutation event by its type, starting position, length, and altered sequence.

### Supplementary Methods

#### Input files to TraceQC R package

A FASTQ file and a reference file are required to use TraceQC. The reference is a text file which contains information as follows:

```
ATGGACTATCATATGCTTACCG...CCGGTAGACGCACCTCCACCCCACAGTGGGGTTAGAGCTAGAAATA
target 23 140
```

The first line of the reference file represents a construct sequence. The second line indicates target region of the construct. In the lines, two numbers next to a region name specify the start and end locations of the region. Locations should be 0-based, i.e. the first location is indicated as 0. If users want to add additional regions like spacer region or PAM region, users can add more lines that contains the additional regions. The format of the regions is the same as the target region. Here is an example of the refenence file with additional regions:

```
ATGGACTATCATATGCTTACCG...CCGGTAGACGCACCTCCACCCCACAGTGGGGTTAGAGCTAGAAATA
target 23 140
spacer 87 107
PAM 107 110
```

#### Main steps in TraceQC

##### Aligning sequence reads to the reference sequence

To align the target sequence with construct reference sequence, TraceQC uses global alignment with affine penalty as implemented in Biopython (Cock et al. 2009). The default match score, mismatch score, gap opening penalty and gap extension penalty is set to 2, $-2$, $-6$ and $-0.1$ respectively. The motivation of choosing a small gap extension penalty is due to the high proportion of indels in CRISPR induced mutations. After the alignment, the adapter regions are trimmed off and the evolving barcode regions are preserved and used to identify mutation events. Sequence-level parallelization using the `multiprocessing` library is applied to speed up the alignment process. The parallelization makes the process about 10 times faster when 16 cores are used.

##### Identification of mutation events

CRISPR induced mutations show great diversity of indels in terms of length and position (Chen et al. 2019). For each sequence read, TraceQC locates every mutation and extracts the follwoing: the mutation type (point mutation, deletion, or insertion), the mutation start position on the reference sequence, the mutation length, and the mutation altered sequence (Supplementary Figure S1-B).

##### Construction of Boolean sequence

In this step, TraceQC aggregates all the sequence reads and identifies $n$ unique mutation events $[m_1, m_2 \dots m_n]$. TraceQC then converts each sequence read into a boolean sequence $B = [b_1, b_2 \dots b_n]$ in which $b_i = TRUE$ means the sequence contains mutation event $m_i$. This Boolean sequence can be directly applied to reconstruct the cell lineage tree.

#### Generating the TraceQC report

In TraceQC package, the `generate_qc_report` function is used to create a QC report. The following script shows how to generate a QC report using the function.

```
library(TraceQC)
obj <- generate_qc_report(
  input_file = system.file("extdata", "test_data",
                           "fastq", "example.fastq", package="TraceQC"),
  ref_file = system.file("extdata", "test_data",
                         "ref", "ref.txt", package="TraceQC"),
  preview = FALSE,
  title = "TraceQC report",
  ncores=1
  )
summary(obj)
```

Once the function has been executed successfully, a report as shown below will be generated. In the example below, we used a sample from (Kalhor, Mali, and Church 2017).

##### TraceQC report

date: 2020-06-05

##### Input files to generate the report

- Input file: `example.fastq`
- Construct file: `ref.txt`

##### Construct structure

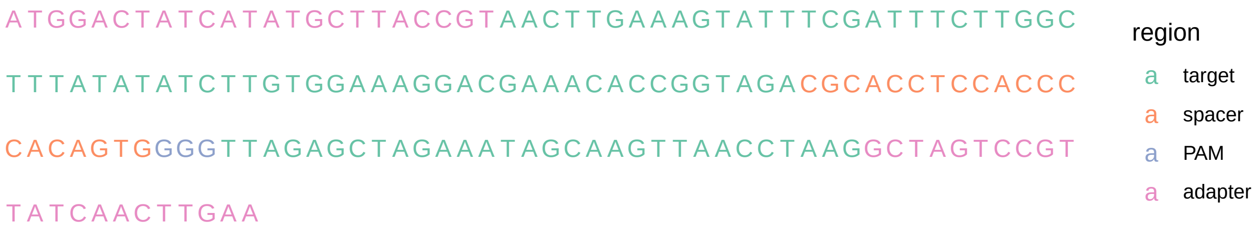

##### Basic Statistics of the sample file

| Measure | Value |
|---|---|
| Filename | example.fastq |
| File type | Conventional base calls |
| Encoding | Sanger / Illumina 1.9 |
| Total Sequences | 5000 |
| Sequences flagged as poor quality | 0 |
| Sequence length | 162 |
| %GC | 41 |

**Sequence quality control**

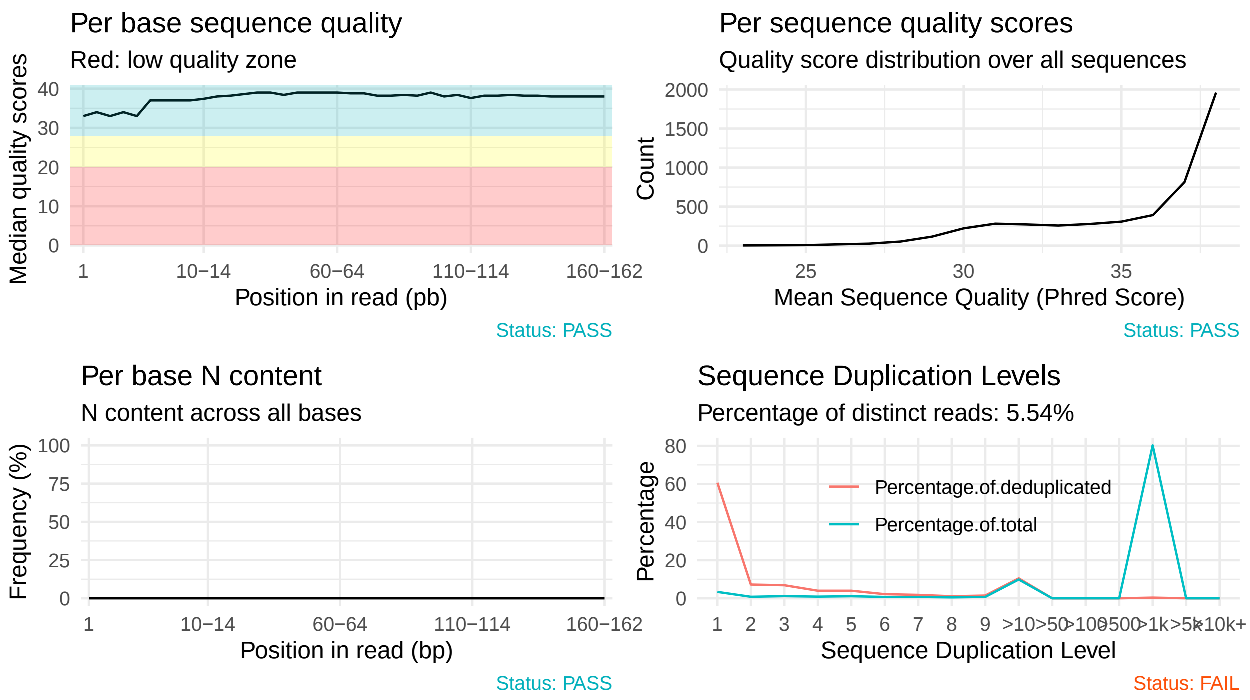

**Alignment score distribution**

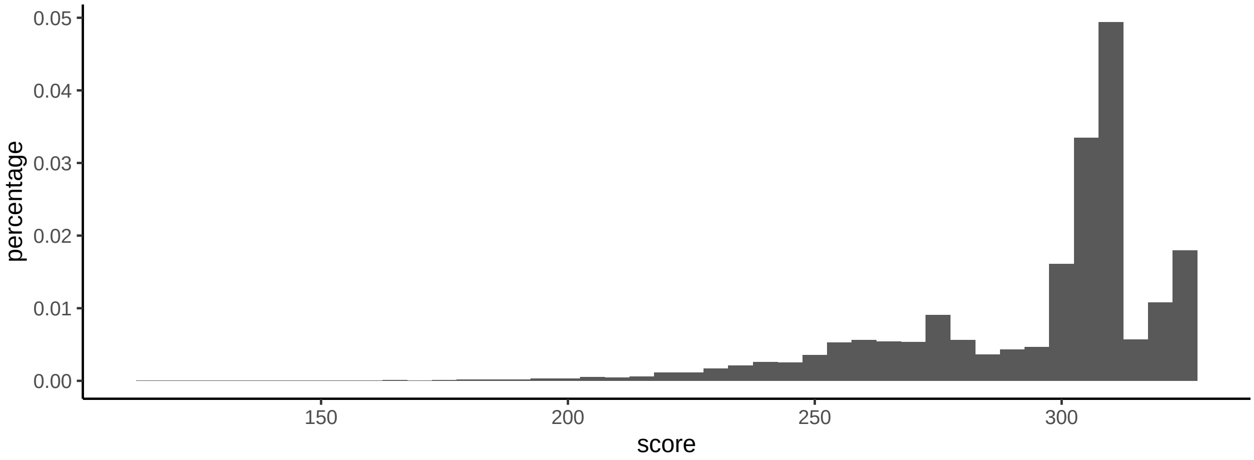

**Barcode distribution inequality**

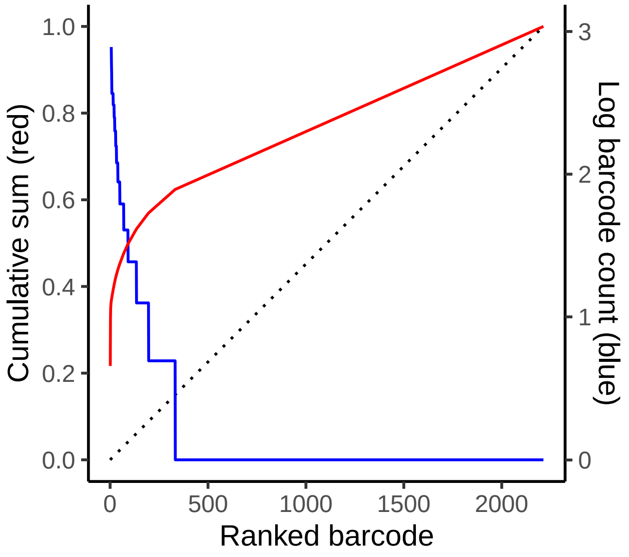

**Most frequent mutation patterns**

| target_seq | type | start | length | mutate_to | count |
|---|---|---|---|---|---|
| ...ACGAAACACCGGTAGACGCACCTCCACCCCA... | insertion | 83 | 1 | A | 1083 |
| ...ACGAAACACCGGTAGACGCACCTCCACCCCA... | unmutated | 0 | 0 | - | 549 |
| ...ACGAAACACCGGTAGACGCACCTCCACCCCA... | insertion | 82 | 2 | AA | 123 |
| ...ACGAAACACCGGTAGACGCACCTCCACCCCA... | deletion | 81 | 14 | - | 31 |
| ...ACGAAACACCGGTAGACGCACCTCCACCC–... | deletion | 79 | 16 | - | 28 |
| ...ACGAAACACCG——————–... | deletion | 61 | 22 | - | 18 |
| ...ACGAAACACCGGTAGACGCACCTCCACC—... | deletion | 78 | 17 | - | 16 |
| ...ACGAAACACCGGTAGACGCAC———–... | deletion | 71 | 24 | - | 15 |
| ...ACGAAACACCGGTAGAC————... | deletion | 67 | 18 | - | 13 |
| ...ACGAAACACCGGTAGACGCACC———... | deletion | 72 | 12 | - | 13 |

**Number of mutations per barcode**

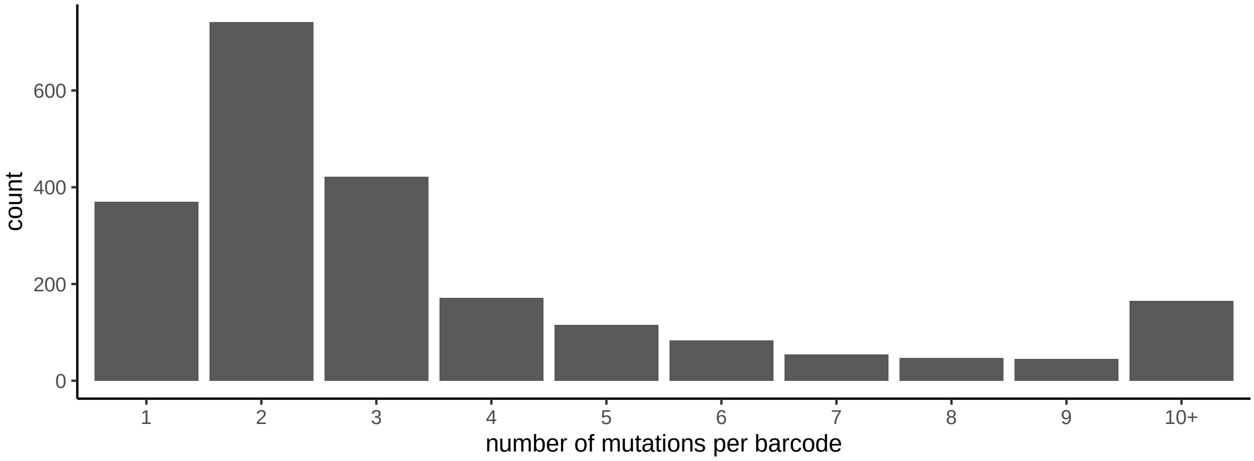

##### Summary of mutation events

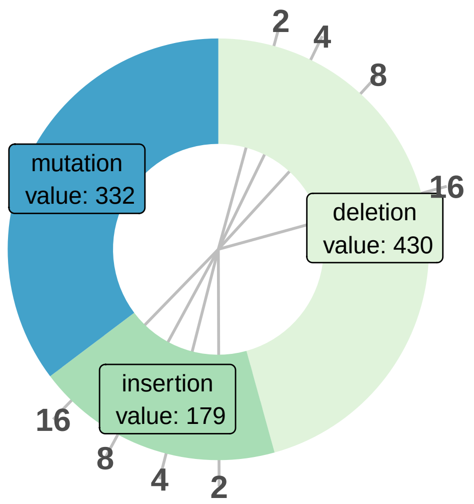

##### Mutation hotspot plots

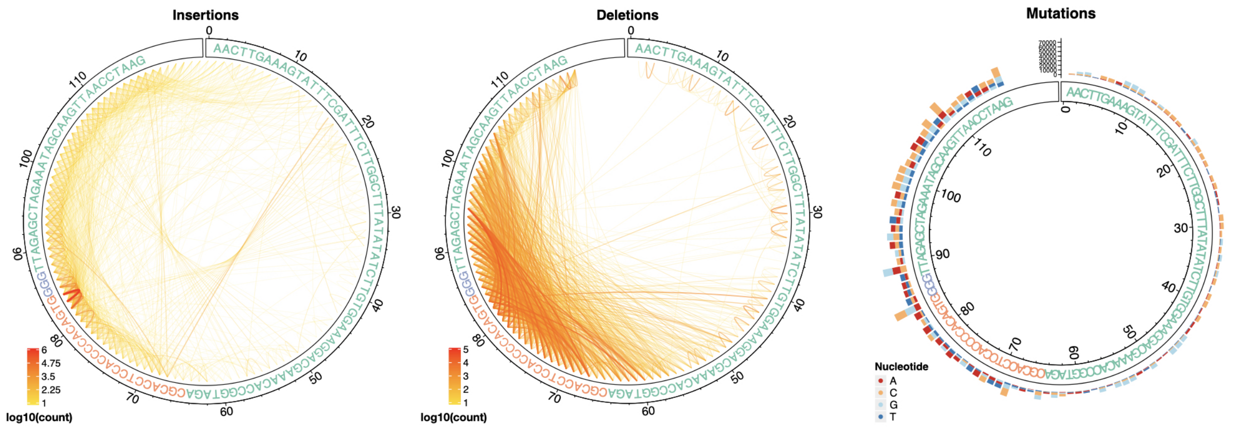

#### Generating a TraceQC object

When users want to use plot functions in TraceQC, it is required to create a TraceQC object for a given sample. This section shows how to create the object.

First, the `TraceQC` package needs to be imported. The package is available at https://github.com/LiuzLab/TraceQC. If there is no FastQC report, it is recommended to import `fastqcr` package to create a FastQC report for the sample.

```
library(TraceQC)
library(fastqcr)
```

To create a TraceQC object, three different files are required.

- `input_file`: A FASTQ file from an experiment of linage tracing experiment using CRISPR.

```
input_file <- system.file("extdata", "test_data",
                          "fastq", "example_small.fastq", package="TraceQC")
cat(readLines(input_file)[1:4], sep = "\n")
```

- `ref_file`: A text file that contains a construct (for reference) sequence.

```
ref_file <- system.file("extdata", "test_data", "ref",
                        "ref.txt", package="TraceQC")
cat(readLines(ref_file), sep = "\n")
```

- `input_qc_path`: A path to the FastQC file which corresponds to `input_file`. It is possible to import the FastQC file from outside the workspace, but if no FastQC file has been generated yet, then it is possible to create it using the `fastqcr` package. The package can be installed by using `install_external_packages`. To generate a FastQC file and get the path, the following lines are needed.

```
qc_dir <- tempdir() # It is possible to set the dir to another location.

# The first argument is a directory, not a path,
# so if there are multiple FASTQ files in a directory, it doesn't have to run
# `fastqc` function multiple times.
fastqc(system.file("extdata", "test_data",
                   "fastq", package = "TraceQC"),
       qc.dir=qc_dir)
# This function tell where the FastQC file which is corresponded to `input_file`.
input_qc_path <- get_qcpath(input_file, qc_dir)
```

After the required files are ready, running `TraceQC` will generate an object.

```
obj <- TraceQC(input_file = input_file,
               ref_file = ref_file,
               fastqc_file = input_qc_path,
               ncores = 4)
```

#### Additional analysis by TraceQC

`TraceQC` is a versatile tool. In addtion to performing Quality Control, it can be used for phylogenetic reconstruction and can handle time series data.

##### Phylogenetic reconstruction

The example below shows how to load an object to run a phylogenetic reconstruction using `phangorn` and `ggtree` package.

First, we are going to load `TraceQC` package and an example object (`example_obj`).

```
library(TraceQC)
library(phangorn)
library(ggtree)
data(example_obj)
```

Next, `build_character_table` in `TraceQC` will convert the object to a list that contains a matrix and sequence information.

```
tree_input <- build_character_table(example_obj)
```

Finally, we can reconstruct a phylogenetic tree with the following code.

```
data <- phyDat(data=tree_input$character_table,type="USER",levels=c(0,1))
dm <- dist.hamming(data)
treeUPGMA <- upgma(dm)
treePars <- optim.parsimony(treeUPGMA, data)
```

```
## Final p-score 68 after  0 nni operations
```

```
ggtree(treePars) +
  geom_tiplab(size=2)
```

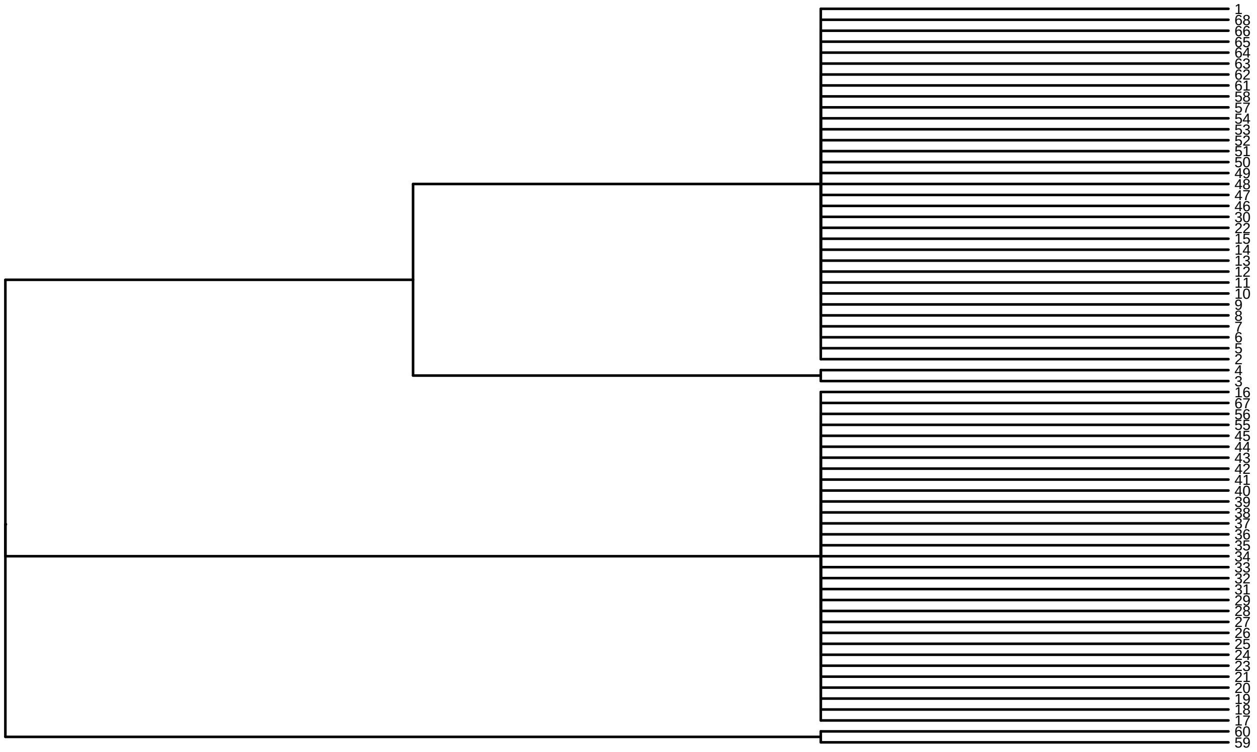
1
68
66
65
64
63
62
61
58
57
54
53
52
51
50
49
48
47
46
30
22
15
14
13
12
11
10
9
8
7
6
5
2
4
3
16
67
56
55
45
44
43
42
41
40
39
38
37
36
35
34
33
32
31
29
28
27
26
25
24
23
21
20
19
18
17
60
59

##### Handling time series data

TraceQC provides a function to handle multiple samples for different time points. The following R script shows how to handle multiple samples using the `create_obj_list` function. In the example below, we use samples of day 0, day 2, and day 14 from (Kalhor, Mali, and Church 2017).

```
samples <- list(
  "day00" = system.file("extdata", "test_data", "fastq",
                        "example_0d.fastq", package="TraceQC"),

  "day02" = system.file("extdata", "test_data", "fastq",
                        "example_2d.fastq", package="TraceQC"),

  "day14" = system.file("extdata", "test_data", "fastq",
                        "example_14d.fastq", package="TraceQC")
)

ref <- system.file("extdata", "test_data", "ref",
                   "ref.txt", package="TraceQC")

obj_list <- create_obj_list(samples, ref)
```

After running `create_obj_list`, `obj_list` which is a list and has three elements is created.

```
summary(obj_list)
```

```
##       Length Class  Mode
## day00 5      -none- list
## day02 5      -none- list
## day14 5      -none- list
```

With obj_list, users can check changes of the percentage of mutations across different time points using plot_mutation_pattern_lineplot or plot_mutation_pattern_violinplot.

```
plot_grid(
  plot_mutation_pattern_lineplot(obj_list),
  plot_mutation_pattern_violinplot(obj_list), ncol=1)
```

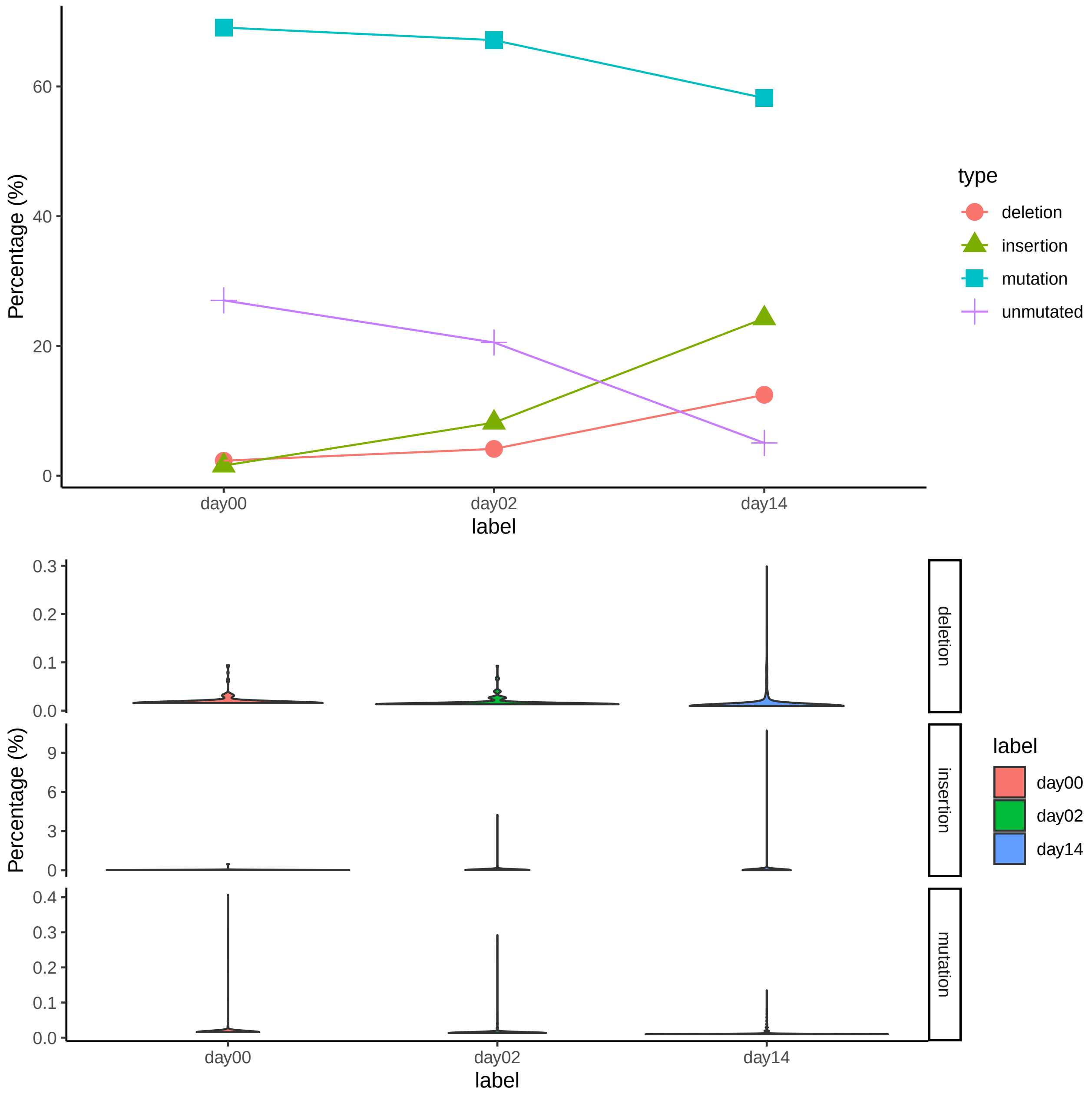

#### Programming libraries

The following programming libraries were used To implement the TraceQC package:

Languages:

- `R` (Team and others 2020)
- `Python` (Van Rossum and Drake Jr 1995)

Packages:

The following `python` packages were used:

- `biopython` (Cock et al. 2009)
- `pandas` (McKinney and others 2011)

The following `R` packages were used:

- `ggplot2` (Wickham 2011)
- `circlize` (Gu et al. 2014)
- `ComplexHeatmap` (Gu, Eils, and Schlesner 2016)

- `tidyverse` (Wickham et al. 2019)
- `fastqcr` ("Fastqcr: Quality Control of Sequencing Data" 2019)
- `rmarkdown` (Xie, Allaire, and Grolemund 2018)
- `kableExtra` (Zhu 2018)
- `RColorBrewer` ("RColorBrewer: ColorBrewer Palettes" 2014)
- `reticulate` ("Reticulate: Interface to 'Python' " 2020)
- `DECIPHER` (Wright 2020)
- `tictoc` ("Tictoc: Functions for Timing R Scripts, as Well as Implementations of Stack and List Structures" 2014)

Van Rossum, Guido, and Fred L Drake Jr. 1995. *Python Reference Manual*. Centrum voor Wiskunde en Informatica Amsterdam.

Wickham, Hadley. 2011. "Ggplot2." *Wiley Interdisciplinary Reviews: Computational Statistics* 3 (2): 180–85.

Wickham, Hadley, Mara Averick, Jennifer Bryan, Winston Chang, Lucy McGowan, Romain François, Garrett Grolemund, et al. 2019. "Welcome to the Tidyverse." *Journal of Open Source Software* 4 (43): 1686.

Wright, Erik. 2020. "DECIPHER: Tools for Curating, Analyzing, and Manipulating Biological Sequences." Bioconductor version: Release (3.11). https://doi.org/10.18129/B9.bioc.DECIPHER.

Xie, Yihui, Joseph J Allaire, and Garrett Grolemund. 2018. *R Markdown: The Definitive Guide*. CRC Press.

Zhu, Hao. 2018. "KableExtra: Construct Complex Table with'kable'and Pipe Syntax." *URL Https://CRAN. R-Project. Org/Package= kableExtra, R Package Version 0.9. 0.*
